# Are tropical reptiles really declining? A six-year survey of snakes in Drake Bay, Costa Rica, and the role of temperature, rain and light

**DOI:** 10.1101/731174

**Authors:** Jose Pablo Barquero-González, Tracie L. Stice, Gianfranco Gómez, Julián Monge-Nájera

## Abstract

**Introduction:** studies in the last two decades have found declining snake populations in both temperate and tropical sites, including informal reports from Drake Bay, Costa Rica.

**Objective:** to investigate if reports of decreasing snake populations in Drake Bay had a real basis, and if environmental factors, particularly temperature, rain and light, have played a role in that decrease.

**Methods:** we worked at Drake Bay from 2012 through 2017 and made over 4000 h of transect counts. Using head flashlights we surveyed a transect covered by lowland tropical rainforest at an altitude of 12–38 m above sea level, near the Agujas River, mostly at 1930–2200 hours. We counted all the snakes that we could see along the transect.

**Results:** snake counts increase from August to September and then decline rapidly. The May snakes/rainfall peaks coincide, but the second snake peak occurs one month before the rain peak; we counted more snakes in dry nights, with the exception of *Imantodes cenchoa* which was equally common despite rain conditions. We saw less *Leptodeira septentrionalis* on bright nights, but all other species were unaffected. Along the six years, the number *of species* with each diet type remained relatively constant, but the number *of individuals* declined sharply for those that feed on amphibians and reptiles. We report *Rhadinella godmani*, a highland species, at 12–38 m of altitude.

**Conclusion:** night field counts of snakes in Drake Bay, Costa Rica, show a strong decline from 2012 through 2017.

---

Snakes are particularly susceptible to population decline because of their long life spans, late sexual maturity, low reproductive frequency, site fidelity and significant mortality among neonates and juveniles (Scott & Seigel, 1992; Shetty & Shine, 2002). They are also good ecological indicators because their populations often reflect fluctuations in the environment and in the populations of their prey (Moore et al., 2003; Madsen, Ujvari, Shine, & Olsson, 2006).

Population status assessments are difficult because of their cryptic life styles and low or sporadic activity (Gibbon et al., 2000), but there have been reports of population decline in temperate regions, including the United States of America (Mount, 1975; Rudolph & Burgdorf, 1997; Conant & Collins, 1998; Hallam, Wheaton, & Fischer, 1998; Tuberville, Bodie, Jensen, LaClaire, & Gibbons, 2000; Winne, Willson, Todd, Andrews, & Gibbons, 2007) and Europe (Reading et al., 2010). The situation may be worse in the tropics, where documented studies are even more scarce (Böhm et al., 2013; Urban, 2015): declines include the extinction of the Round Island Burrowing Boa, endemic to Mauritius (Bullock, 1986; Greene, 2000), and population drops of snakes in Nigeria (Reading et al., 2010) and Australia (Lukoschek, Beger, Ceccarelli, Richards, & Pratchett, 2013; Lukoschek, 2018).

Even though snake declines seem to be a reality in many parts of the world, hundreds of samples are needed to detect real declines (Kery, 2002; Sewell, Guillera-Arroita, Griffiths, & Beebee, 2012; Hileman et al., 2018). Furthermore, problematic assumptions used in mathematical models can lead to suspicious conclusions (e.g. exaggerated extinction rate estimates: McCallum, 2007).

Reports of snake declines are frequently based on anecdotal evidence and there is a need for studies that provide intensive counts for prolonged periods (Krysko, 2001; Böhm et al., 2013; Urban, 2015); for example, local tourist guides in Drake Bay, Costa Rica, have told us that populations have been declining for nearly two decades. Our objective in this study was to investigate if their impression is matched by more formal data collection.

## MATERIALS AND METHODS

We worked at Drake Bay, South Pacific of Costa Rica (N 08.69420–08.69490; W 083.67421–083.67495), from 2012 through 2017, and made over 4 000 h of transect counts of snakes. These counts were made while we were accompanied by small groups of tourists as part of our work as field guides in the area.

We hypothesized that climate changes may reduce local populations of some species and increase others; and that snakes that feed exclusively on amphibians or insect prey (which fluctuate strongly with rain or temperature), should be more affected than generalist species.

### Snake counts

Using head flashlights we surveyed a transect covered by lowland tropical rainforest, 12–38 m above sea level, near the Agujas River (Fig. 1 in Digital Appendix 1). We counted all the snakes that we could see along the transect; identified them *in situ* (guides by Savage, 2002 and Solórzano, 2004), and photographed them for taxonomic corroboration (Digital Appendix 2). We worked mostly from 1930 to 2200 hours, but a few counts started at 1730 and ended at 2245 hours (sampling hours per date in Digital Appendix 2).

### Precipitation rates and temperature

Precipitation rates and temperature data were kindly provided by the *Instituto Meteorológico Nacional de Costa Rica* from the nearest meteorological station at Rancho Quemado.

#### Note

Rancho Quemado, where the meteorological station is located, has an altitude of 240 m above sea level while the survey site in Drake Bay has an altitude of 12 to 38 m, so we expected a difference in temperature between the study site and the meteorological station. To assess this difference, we made temperature measurements directly in Drake Bay from February 2017 to August 2017 and compared them with Rancho Quemado for that same time period. We found that average temperatures in Drake Bay are 2 to 3 degrees Celsius higher than temperatures in Rancho Quemado, but that *the trends along the timeline are the same*. No precipitation records were available from January 2012 to July 2012, December 2016 to March 2017 and September 2017 to December 2017. No temperature records were available from January 2012 to July 2012 and from September 2017 to December 2017.

### Moon and *in situ* rain

Instead of using published moon phase data, we recorded moonlight conditions on every trip directly in the trail because cloudy skies produce dark nights even when the moon is full. We used both official rain records and our own classification of rain condition during the trip (see Results).

### Statistical analyses

We analyzed the counts independently for each of the most common species, and pooled the rare species into an “others” category that we also analyzed (Table 2 and Table 3 in Digital Appendix 1).

### Ethical, conflict of interest and financial statements

the authors declare that they have fully complied with all pertinent ethical and legal requirements, both during the study and in the production of the manuscript; that there are no conflicts of interest of any kind; that all financial sources are fully and clearly stated in the acknowledgements section; and that they fully agree with the final edited version of the article. Tracie L. Stice and Gianfranco Gómez collected the data, Jose Pablo Barquero-González and Julián Monge-Nájera analyzed the data, all the authors participated in writing the manuscript. We did not need collecting permits because we did not collect any animals. A signed document with this statement has been filed in the journal archives.

## RESULTS

### Species observed

In total, we recorded 25 snake species, representing five families (Boidae, Colubridae, Dipsadidae, Elapidae, Viperidae) (Table 1 in Digital Appendix 1).

### Effect of diet

Along the six years, the number of species with each diet type remained relatively constant, fluctuating by a couple of species each year (Fig. 2 in Digital Appendix 1). However, the numbers of individuals changed visibly along the six-year study period depending on their main diet items. Species that mostly eat fish and invertebrates were always rare, and thus insufficient for us to see temporal trends; snakes that mostly feed on birds and mammals, which had larger populations, declined from 2012 through 2015. Finally, species that feed mostly on reptiles and amphibians were initially abundant but also have constantly declined over the six-year period, despite some occasional population peaks (Fig. 3 in Digital Appendix 1).

### Annual patterns

When we compared counts of the five most frequent species with rainfall and temperature during the six-year study period, we noticed several trends (Fig. 4 in Digital Appendix 1). One is a weak increase in overall temperature along the study period. The other is that numerically dominant species have highly specific patterns but most started a decline since 2015. We observed a decrease in *L. septentrionalis* and *I. cenchoa* since 2015; *E. sclateri* is scarce (except for a peak from August to September of 2012 and 2014) but it has become even rarer since 2015; *S. compresus* was not seen after April 2016; finally, *M. melanolomus* had no clear tendency to grow or decline from 2012 to 2017, but showed a peak between August to September (Fig. 4 in Digital Appendix 1).

### Monthly pattern

Mean temperature does not change strongly along the year, but rainfall increases after February and reaches a maximum in October, while snake counts increase from August to September and then decline rapidly. The May snakes/rainfall peaks coincide, but the second snake peak occurs one month before the rain peak (Fig. 5 in Digital Appendix 1).

### Ecological correlations

The statistical significance values of ANOVA tests appear in Table 2 and Table 3 (Digital Appendix 1) and the counts/environmental condition relationships in the appendices,

### Effect of rain

We counted more snakes in dry nights, with the exception of *I. cenchoa* which was equally common despite rain conditions (Fig. 6 in Digital Appendix 1).

### Effect of moonlight

We saw less *L. septentrionalis* on bright nights, but all other species were unaffected so they are not presented in the figure (Fig. 7 in Digital Appendix 1).

### Final considerations

We did not see differences in leaf litter quantity from the beginning to the end of the study period (Digital Appendix 2). Our finding of *R. godmani* is unexpected because it is a highland species (see Discussion).

## DISCUSSION

### Food

Snake activity patterns depend on changes in food availability throughout the year (Henderson, Dixon, & Soini, 1978; Martins, 1994). Snake species that ambush their prey and rely on sit-and-wait strategies are particularly susceptible because of their low rate of food acquisition (Webb & Shine, 1998), just like specialist species, which are less likely to exploit alternative resources in response to shifting environmental conditions (Terborgh & Winter, 1980; Gaston, 1994). Counts of *I. cenchoa*, a snake that feeds mainly on reptiles, sharply fell in 2015 and 2017; *E. sclateri*, a specialist feeder on reptile eggs, declined in 2015 and almost disappeared in 2016–2017, possibly due to scarce prey. The fall in the numbers of *L. septentrionalis* matches the fall in the abundance of prey species like *Boana rosenbergi, Smilisca phaeota* and *Agalychnis callidryas*; nevertheless, other prey items for this species, like *Rhinella horribilis* and *Craugastor fitizingeri*, remained common (Gianfranco Gómez, personal observation).

If we consider that four of the five species affected by rainfall in our study have diets consisting primarily of amphibians, reptiles, or both, our results are consistent with those of a study in Mexico, where more rain lead to more amphibian activity and to increased populations of the snakes that feed on them (Duellman, 1958).

*Rhadinella godmani* is a small leaf litter snake of uncertain diet and considered a highland species (Savage, 2002; Solórzano, 2004); we ignore if its presence at near sea level in Drake is in any way related to changes in environmental factors.

### Temperature

Higher temperatures reduce the metabolic rate of snakes and restrain their activity and visibility (Seigel, Collins, & Novak, 1987; Zamora-Camacho, Moreno-Rueda, & Pleguezuelos, 2010; Rugiero, Milana, Petrozzi, Capula, & Luiselli, 2013). In Drake Bay, average monthly temperatures did not vary widely, so we are not surprised that there were no strong, generalized trends in species abundance that could be related with temperature. This matches previous work with tropical species (e.g. Shine & Madsen, 1996; Luiselli & Akani, 2002). Only one species, *S. compressus*, which needs fresh, shady habitats, was not recorded at all from our sampling site along 2016 and 2017.

### Rain

There are reports of no correlation between counts of Neotropical snakes and rainfall (Henderson & Hoevers, 1977; Martins, 1994; Bernarde & Abe, 2006) and this is similar to our finding that the arboreal *I. cenchoa* was equally common despite rain conditions. Other species become more active with rain (Daltry, Ross, Thorpe, & Wüster, 1998; Oliveira & Martins, 2001; Morrison & Bolger, 2002), but we did not see any species more often in rainy nights in Drake, quite the opposite, Drake snake counts were higher in dry nights. Perhaps they avoid the cold rainwater (rain is cold in the tropics, too).

### Moonlight

Snakes may increase activity in full moon nights, when prey are more visible (Lillywhite & Brischoux, 2012; Connoly & Orrock, 2018); for example, the tropical tree snake *Boiga irregularis* even moves to areas where moonlight is stronger (Campbell, Mackessy, & Clarke, 2008). However, *Crotalus viridis* avoids bright moonlight, possibly to escape detection by its predators (Clarke, Chopko, & Mackessy, 1996). In our study, the fewer sightings of *L. septentrionalis* on nights with strong moonlight may also mean that it is avoiding predation, but we saw all the other species at Drake with the same frequency in dark and illuminated nights.

### Final remarks

The clear decline in snake counts at Drake along the years might mean that their populations have decreased, that they moved out of sight or that they migrated to higher, cooler areas. They did not seem to avoid the transect because of our presence there (actually a few species remained constant in our counts along the years) and we do not think they found a suitable habitat at higher altitude because of human alteration of habitats around the reserve. We believe that the population decline in Drake is real and needs attention from the conservation authorities.

## Supporting information

Digital appendix

## ACKNOWLEDGMENTS

We thank Carolina Seas for her assistance with data analysis, Sergio Aguilar for his help in the elaboration of the images, Alejandro Solórzano, Héctor Zumbado and Mahmood Sasa for recommendations to improve the manuscript, and Instituto Meteorológico Nacional for climatic data. This study was financed by the authors.

## RESUMEN

### ¿Están disminuyendo los reptiles tropicales? Seis años de monitoreo en las serpientes de Bahía Drake, Costa Rica, y el papel de la temperatura, la lluvia y la luz

#### Introducción

los estudios realizados en las últimas dos décadas han encontrado una disminución de las poblaciones de serpientes en ecosistemas templados y tropicales, incluyendo observaciones informales en Bahía Drake, Costa Rica.

#### Objetivo

investigar si los informes de disminución de las poblaciones de las serpientes de Bahía Drake tuvieron una base real, y si los factores ambientales, particularmente temperatura, lluvia y luz, han jugado un papel en esa disminución.

#### Métodos

trabajamos en Bahía Drake desde el 2012 hasta el 2017 y realizamos más de 4 000 h de recuentos de serpientes. Usando linternas de cabeza y en un horario mayormente entre las 1930 y las 2200 horas, examinamos un transecto de bosque tropical de bajura a 12–38 m.s.n.m, cerca del río Agujas.

#### Resultados

la cantidad de serpientes vistas aumenta de agosto a setiembre y luego disminuye rápidamente. Los picos de serpientes y lluvia coinciden en mayo, pero el segundo pico de serpientes es un mes antes del pico de lluvia; contamos más serpientes en las noches secas, con la excepción de *Imantodes cenchoa* que era igualmente común en todas las condiciones de lluvia. Vimos menos *Leptodeira septentrionalis* en noches brillantes, pero las demás especies no se vieron afectadas por la luz lunar. A lo largo de los seis años, la *cantidad de especies* con cada tipo de dieta se mantuvo relativamente constante, pero la *cantidad de individuos* disminuyó considerablemente en las especies que se alimentan de anfibios y reptiles. Hallamos *Rhadinella godmani*, una especie de montaña, a 12–38 m de altitud.

#### Conclusión

los conteos nocturnos de serpientes en Bahía Drake, Costa Rica, muestran una fuerte disminución del 2012 al 2017.

#### Palabras clave

demografía de serpientes, luz de la luna, lluvia, temperature, cambio climático en Osa.

## REFERENCIAS

Bernarde, P. S., & Abe, A. S. (2006). A snake community at Espigão do Oeste, Rondônia, southwestern Amazon, Brazil. South American Journal of Herpetology, 1(2),102–113. DOI: 10.2994/1808-9798(2006)1[102:ASCAED]2.0.CO;2

Böhm, M., Collen, B., Baillie, J. E., Bowles, P., Chanson, J., Cox, N., & Rhodin. A. G. (2013). The conservation status of the world’s reptiles. Biological Conservation, 157, 372–385. DOI: 10.1016/j.biocon.2012.07.015

Bullock, D. J. (1986). The ecology and conservation of reptiles on Round Island and Gunner’s Quion, Mauritius. Biological Conservation, 37(2) 135–156. DOI: 10.1016/0006-3207(86)90088-1

Campbell, S. R., Mackessy, S. P., & Clarke, J. A. (2008). Microhabitat use by brown treesnakes (Boiga irregularis): effects of moonlight and prey. Journal of Herpetology, 42(2), 246-250. DOI: 10.1655/07-054.1

Clarke, J. A., Chopko, J. T., & Mackessy, S. P. (1996). The effect of moonlight on activity patterns of adult and juvenile prairie rattlesnakes (*Crotalus viridis viridis*). Journal of Herpetology, 30(2), 192–197. DOI: 10.2307/1565509

Conant, R., & Collins, J. T. (1998). Reptiles and Amphibians of North America. 4th Edition. New York, USA: Houghton Mifflin.

Connolly, B. M., & Orrock, J. L. (2018). Habitat-specific capture timing of deer mice (*Peromyscus maniculatus*) suggests that predators structure temporal activity of prey. Ethology, 124(2), 105–112. DOI: 10.1111/eth.12708

Daltry, J. C., Ross, T., Thorpe, R. S., & Wüster, W. (1998). Evidence that humidity influences snake activity patterns: a field study of the Malayan pit viper *Calloselasma rhodostoma*. Ecography, 21(1), 25–34. DOI: 10.1111/j.1600-0587.1998.tb00391.x

Duellman, W. E. (1958). A monographic study of the colubrid snake genus *Leptodeira*. Bulletin of the American Museum of Natural History, 114, article 1.

Gaston, K. J. (1994). What is rarity? In K. J. Gaston (Ed.). Rarity (pp. 1–21). Dordrecht, Netherlands: Springer. DOI: 10.1007/978-94-011-0701-3_1

Gibbons, J. W., Scott, D. E., Ryan, T. J., Buhlmann, K. A., Tuberville, T. D., Metts, B. S., … & Winne, C. T. (2000). The Global Decline of Reptiles, Déjà Vu Amphibians. BioScience, 50(8), 653–666. DOI: 10.2307/1445695

Greene, H. W. (2000). Snakes: the evolution of mystery in nature. Los Angeles, CA, USA.: University of California Press.

Hallam, C. O., Wheaton, K., & Fischer R. A. (1998). *Species Profile: Eastern Indigo Snake (Drymarchon corals couperi) on Military Installations in the Southeastern United States* (No. WES-TR-SERDP-98-2). Technical Report, Army engineer waterways experiment station Vicksburg, MS, USA. DOI: 10.21236/ADA342329

Henderson, R. W., & Hoevers, L.G. (1977). The seasonal incidence of snakes at a locality in northern Belize. Copeia, 1977, 349–355. DOI: 10.2307/1443914

Henderson, R. W., Dixon, J. R., & Soini, P. (1978). On the seasonal incidence of tropical snakes. Wisconsin, USA: Milwaukee Public Museum.

Hileman, E. T., Allender, M. C., Bradke, D. R, Faust, L. J., Moore, J. A., Ravesi, M. J., & Tetzlaff. S. J. (2018). Estimation of Ophidiomyces prevalence to evaluate snake fungal disease risk. The Journal of Wildlife Management, 82(1), 173–181. DOI: 10.1002/jwmg.21345

Kery, M. (2002). Inferring the absence of a species: a case study of snakes. The Journal of wildlife management, 66(2), 330–338. DOI: 10.2307/3803165

Krysko, K. L. (2001). Ecology, conservation, and morphological and molecular systematics of the kingsnake, Lampropeltis getula (Serpentes: Colubridae). (Ph.D Dissertation). University of Florida, Florida, USA.

Lillywhite, H. B., & Brischoux, F. (2012). Is it better in the moonlight? Nocturnal activity of insular cottonmouth snakes increases with lunar light levels. Journal of Zoology, 286(3), 194–199. DOI: 10.1111/j.1469-7998.2011.00866.x

Lukoschek, V., Beger, M., Ceccarelli, D., Richards, Z., & Pratchett, M. (2013). Enigmatic declines of Australia’s sea snakes from a biodiversity hotspot. Biological Conservation, 166, 191–202. DOI: 10.1016/j.biocon.2013.07.004

Lukoschek, V. (2018). Population declines, genetic bottlenecks and potential hybridization in sea snakes on Australia’s Timor Sea reefs. Biological Conservation, 225, 66–79. DOI: 10.1016/j.biocon.2018.06.018

Luiselli, L., & Akani, G. C. (2002). Is thermoregulation really unimportant for tropical reptiles? Comparative study of four sympatric snake species from Africa. Acta Oecologica, 23(2), 59–68. DOI: 10.1016/S1146-609X(02)01134-7

Madsen, T., Ujvari, B., Shine, R., & Olsson, M. (2006). Rain, rats and pythons: Climate-driven population dynamics of predators and prey in tropical Australia. Austral Ecology, 31(1), 30–37. DOI: 10.1111/j.1442-9993.2006.01540.x

Martins, M. (1994). História natural e ecologia de uma taxocenose de serpentes de mata na região de Manaus, Amazônia Central, Brasil. Campinas, SP. (Ph.D Dissertation). Universidade Estadual de Campinas, Brasil.

McCallum, M. L. (2007). Amphibian decline or extinction? Current declines dwarf background extinction rate. Journal of Herpetology, 41(3), 483–491. DOI: 10.1670/0022-1511(2007)41[483:ADOECD]2.0.CO;2

Moore, J. L., Balmford, A., Brooks, T., Burgess, N. D., Hansen, L. A., Rahbek, C., & Williams, P. H. (2003). Performance of sub-Saharan vertebrates as indicator groups for identifying priority areas for conservation. Conservation Biology, 17(1), 207–218. DOI: 10.1046/j.1523-1739.2003.01126.x

Morrison, S. A., & Bolger, D. T. (2002). Variation in a sparrow’s reproductive success with rainfall: food and predator-mediated processes. Oecologia, 133(3), 315–324. DOI: 10.1007/s00442-002-1040-3

Mount, R. H. (1975). The Reptiles and Amphibians of Alabama. Auburn (AL), USA: Auburn University, Alabama Agricultural Experimental Station.

Oliveira, M. E., & Martins, M. (2001). When and where to find a pitviper: activity patterns and habitat use of the lancehead, *Bothrops atrox*, in central Amazonia, Brazil. Herpetological Natural History, 8(2), 101–110.

Reading, C. J., Luiselli, L. M., Akani, G. C., Bonnet, X., Amori, G., Ballouard, J. M., … & Rugiero, L. (2010). Are snake populations in widespread decline? Biology letters, 6(6), 777–780. DOI: 10.1098/rsbl.2010.0373

Rudolph, D. C., & Burgdorf, S. J. (1997). Timber rattlesnakes and Louisiana pine snakes of the west Gulf Coastal Plain: hypotheses of decline. Texas Journal of Science, 49(3), 111–122.

Rugiero, L., Milana G., Petrozzi, F., Capula, M., & Luiselli, L. (2013). Climate-change-related shifts in annual phenology of a temperate snake during the last 20 years. Acta oecologica, 51, 42–48. DOI: 10.1016/j.actao.2013.05.005

Scott, N. J. Jr., & Seigel, R. A. (1992). The management of amphibians and reptile populations: Specific priorities and methodological and theoretical constraints. In D. R. McCullough, & R. H. Barrett (Eds.). Wildlife 2001: Populations (pp. 343–368). London, England: Elsevier Applied Science. DOI: 10.1007/978-94-011-2868-1_29

Savage, J. M. (2002). The amphibians and reptiles of Costa Rica: a herpetofauna between two continents, between two seas. Chicago, Illinois, USA: University of Chicago.

Seigel, R. A., Collins, J. T., & Novak, S. S. (1987). Snakes: ecology and evolutionary biology, New York, USA: MacMillan Publishing Company. DOI: 10.2307/1445695

Sewell, D., Guillera-Arroita, G., Griffiths, R. A., & Beebee, T. J. (2012). When is a species declining? Optimizing survey effort to detect population changes in reptiles. PloS one, 7(8), e43387. DOI: 10.1371/journal.pone.0043387

Shetty, S., & Shine, R. (2002). Philopatry and homing behavior of sea snakes (*Laticauda colubrina*) from two adjacent islands in Fiji. Conservation Biology, 16(5), 1422–1426. DOI: 10.1046/j.1523-1739.2002.00515.x

Shine, R., & Madsen, T. (1996). Is thermoregulation unimportant for most reptiles? An example using water pythons (*Liasis fuscus*) in tropical Australia. Physiological Zoology, 69(2), 252–269. DOI: 10.1086/physzool.69.2.30164182

Solórzano, A. (2004). Serpientes de Costa Rica: distribución, taxonomía e historia natural. San José, Costa Rica: Editorial INBio.

Terborgh, J., & Winter, B. (1980). Some causes of extinction. Conservation biology: an evolutionary-ecological perspective. Sinauer Associates, Sunderland, Massachusetts, 1980, 119–133.

Tuberville, T. D., Bodie, J. R., Jensen, J. B., LaClaire, L., & Gibbons, J. W. (2000). Apparent decline of the southern hog-nosed snake, *Heterodon simus*. Journal of the Elisha Mitchell Scientific Society, 116(1), 19–40.

Urban, M. C. (2015). Accelerating extinction risk from climate change. Science, 348(6234), 571–573. DOI: 10.1126/science.aaa4984

Webb, J. K., & Shine, R. (1998). Ecological characteristics of a threatened snake species, *Hoplocephalus bungaroides* (Serpentes, Elapidae). Animal Conservation, 1(3), 185–193. DOI: 10.1111/j.1469-1795.1998.tb00028.x

Winne, C. T., Willson, J. D., Todd, B. D., Andrews, K. M., & Gibbons, J.W. (2007). Enigmatic decline of a protected population of Eastern Kingsnakes, *Lampropeltis getula*, in South Carolina. Copeia, 2007(3), 507–519. DOI: 10.1643/0045-8511(2007)2007[507:EDOAPP]2.0.CO;2

Zamora-Camacho, F. J., Moreno-Rueda, G., & Pleguezuelos, J. M. (2010). Long-and short-term impact of temperature on snake detection in the wild: further evidence from the snake Hemorrhois hippocrepis. Acta Herpetologica, 5(2), 143–150.

