## Supplementary material for "Are tropical reptiles really declining? A six-year survey of snakes in Drake Bay, Costa Rica, and the role of temperature, rain and light": Digital appendix: DIGITAL APPENDIX 1.docx

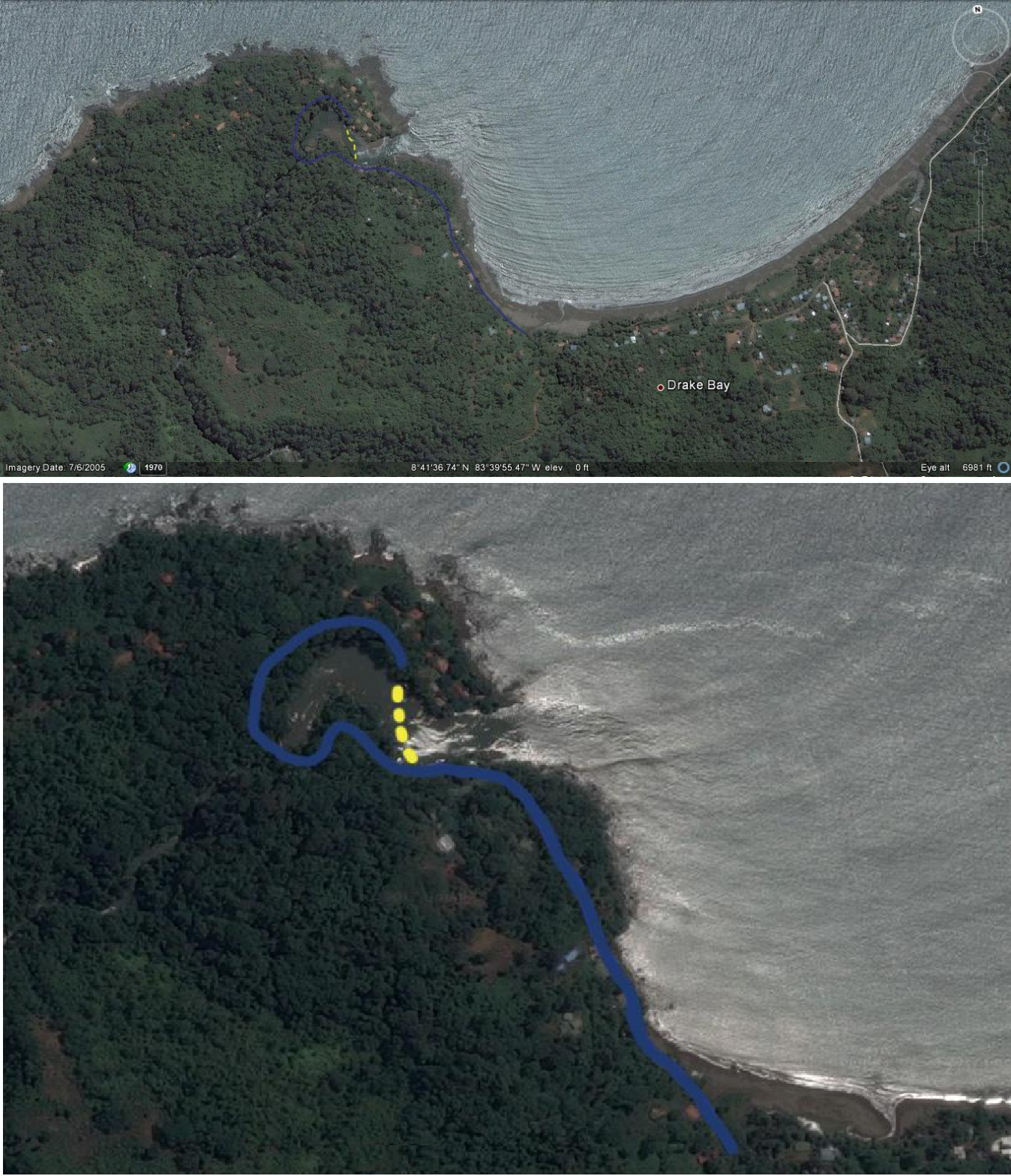

**Fig. 1**. Sampled transect where we counted snakes near the Agujas River, Drake Bay, Costa Rica (lower image: detail of area in upper image)- Aerial photographs from Google Earth; see Methods for coordinates).

TABLE 1

Snake species found on the surveyed years and their main prey items according to the literature (Savage, 2002; Solórzano, 2004)).

| **Prey item** | | | | | | |
| --- | --- | --- | --- | --- | --- | --- |
| **Species** | **Invertebrates** | **Fish** | **Amphibians** | **Reptiles** | **Birds** | **Mammals** |
| *Boa imperator* | - | - | - | X | X | X |
| *Bothriechis schlegelii* | - | - | X | X | X | X |
| *Bothrops asper* | - | - | X | X | X | X |
| *Chironius flavopictus* | - | - | X | - | - | - |
| *Clelia clelia* | - | - | - | X | X | X |
| *Coniophanes fissidens* | - | - | X | X |  | - |
| *Corallus ruschenbergerii* | - | - | - | X | X | X |
| *Dendrophidion percarinatum* | - | - | X | - | - | - |
| *Enuliophis sclateri* | - | - | - | X | - | - |
| *Hydrophis platurus* | - | X | - | - | - | - |
| *Imantodes cenchoa* | - | - | X | X | - | - |
| *Leptodeira septentrionalis* | - | - | X | X | - | - |
| *Leptophis ahaetulla* | - | - | X | X | X | - |
| *Leptophis nebulosus* | - | - | X | - | - | - |
| *Mastigodryas melanolomus* | - | - | X | X | X | X |
| *Micrurus alleni* | - | - | X | X | - | - |
| *Ninia maculata* | X | - | - | - | - | - |
| *Oxyrhopus petolarius* | - | - | X | X | - | X |
| *Phrynonax poecilonotus* | - | - | - | - | X | X |
| *Rhadinella godmani* | - | - | X | X | - | - |
| *Sibon nebulatus* | X | - | - | - | - | - |
| *Siphlophis compressus* | - | - | - | X | - | - |
| *Spilotes pullatus* | - | - | - | X | X | X |
| *Stenorrhina degenhardtii* | X | - | - | - | - | - |
| *Tantilla supracincta* | X | - | - | - | - | - |

TABLE 2

ANOVA results for *Rainfall* versus *Counts per hour* of the most common snake species, Drake Bay, Costa Rica.

| ***Leptodeira septentrionalis.*** | | | | | | | | | | | | |
| --- | --- | --- | --- | --- | --- | --- | --- | --- | --- | --- | --- | --- |
| Variance analysis | | | | | | | | | | | | |
| Variable | | N | | | R^2^ | | | R^2^ Aj | | | CV | |
| Rate | | 216 | | | 0,03 | | | 0,02 | | | 131,61 | |
| Variance analysis table (SC type III) | | | | | | | | | | | | |
| F.V. | SC | | | gl | | | CM | | F | | | p-value |
| Model | 0,25 | | | 2 | | | 0,13 | | 3,08 | | | 0,0482 |
| Weather | 0,25 | | | 2 | | | 0,13 | | 3,08 | | | 0,0482 |
| Error | 8,75 | | | 213 | | | 0,04 | |  | | |  |
| Total | 9,00 | | | 215 | | |  | |  | | |  |
| Adjusted median, standard error and number of observations  Error: 0,0411 df: 213 | | | | | | | | | | | | |
| Weather | | | Median | | | n | | | | E.E. | | |
| Rainy | | | 0,12 | | | 72 | | | | 0,02 | | |
| Light Rain | | | 0,14 | | | 72 | | | | 0,02 | | |
| Dry | | | 0,20 | | | 72 | | | | 0,02 | | |

| ***Imantodes cenchoa.*** | | | | | | | | | | | | |
| --- | --- | --- | --- | --- | --- | --- | --- | --- | --- | --- | --- | --- |
| Variance analysis | | | | | | | | | | | | |
| Variable | | N | | | R^2^ | | | R^2^ Aj | | | CV | |
| Rate | | 216 | | | 0,02 | | | 0,01 | | | 165,09 | |
| Variance analysis table (SC type III) | | | | | | | | | | | | |
| F.V. | SC | | | gl | | | CM | | F | | | p-value |
| Model | 0,04 | | | 2 | | | 0,02 | | 2,11 | | | 0,1233 |
| Weather | 0,04 | | | 2 | | | 0,02 | | 2,11 | | | 0,1233 |
| Error | 1,81 | | | 213 | | | 0,01 | |  | | |  |
| Total | 1,85 | | | 215 | | |  | |  | | |  |
| Test: Tukey Alfa=0,05 DMS=0,03606  Error: 0,0085 df: 213 | | | | | | | | | | | | |
| Weather | | | Median | | | n | | | | E.E. | | |
| Rainy | | | 0,05 | | | 72 | | | | 0,01 A | | |
| Light Rain | | | 0,05 | | | 72 | | | | 0,01 A | | |
| Dry | | | 0,07 | | | 72 | | | | 0,01 A | | |
| ***Enuliophis sclateri.*** | | | | | | | | | | | | |
| Variance analysis | | | | | | | | | | | | |
| Variable | | N | | | R^2^ | | | R^2^ Aj | | | CV | |
| Rate | | 216 | | | 0,07 | | | 0,06 | | | 246,90 | |
| Variance analysis table (SC type III) | | | | | | | | | | | | |
| F.V. | SC | | | gl | | | CM | | F | | | p-value |
| Model | 0,04 | | | 2 | | | 0,02 | | 8,30 | | | 0,0003 |
| Weather | 0,04 | | | 2 | | | 0,02 | | 8,30 | | | 0,0003 |
| Error | 0,48 | | | 213 | | | 2, 3E-03 | |  | | |  |
| Total | 0,52 | | | 215 | | |  | |  | | |  |
| Test: Tukey Alfa=0,05 DMS=0,01866  Error: 0,0023 df: 213 | | | | | | | | | | | | |
| Weather | | | Median | | | N | | | | E.E. | | |
| Rainy | | | 0,01 | | | 72 | | | | 0,01 A | | |
| Light Rain | | | 0,02 | | | 72 | | | | 0,01 A | | |
| Dry | | | 0,04 | | | 72 | | | | 0,01 B | | |
| **Other species (all data from less common species pooled)*.*** | | | | | | | | | | | | |
| Variance analysis | | | | | | | | | | | | |
| Variable | | N | | | R^2^ | | | R^2^ Aj | | | CV | |
| Rate | | 216 | | | 0,03 | | | 0,02 | | | 281,14 | |
| Variance analysis table (SC type III) | | | | | | | | | | | | |
| F.V. | SC | | | gl | | | CM | | F | | | p-value |
| Model | 0,23 | | | 2 | | | 0,11 | | 3,20 | | | 0,0429 |
| Weather | 0,23 | | | 2 | | | 0,11 | | 3,20 | | | 0,0429 |
| Error | 7,63 | | | 213 | | | 0,04 | |  | | |  |
| Total | 7,86 | | | 215 | | |  | |  | | |  |
| Test: Tukey Alfa=0,05 DMS=0,07404  Error: 0,0358 df: 213 | | | | | | | | | | | | |
| Weather | | | Median | | | N | | | | E.E. | | |
| Rainy | | | 0,03 | | | 72 | | | | 0,02 A | | |
| Light Rain | | | 0,07 | | | 72 | | | | 0,02 A B | | |
| Dry | | | 0,11 | | | 72 | | | | 0,02 B | | |

TABLE 3

ANOVA results for *Moonlight* versus *Counts per hour* of snakes, Drake Bay, Costa Rica.

| ***Leptodeira septentrionalis.*** | | | | | | | | | | | | |
| --- | --- | --- | --- | --- | --- | --- | --- | --- | --- | --- | --- | --- |
| Variance analysis | | | | | | | | | | | | |
| Variable | | N | | | R^2^ | | | R^2^ Aj | | | CV | |
| Rate | | 216 | | | 0,04 | | | 0,03 | | | 116,75 | |
| Variance analysis table (SC type III) | | | | | | | | | | | | |
| F.V. | SC | | | gl | | | CM | | F | | | p-value |
| Model | 0,33 | | | 2 | | | 0,17 | | 4,70 | | | 0,0100 |
| Moonlight | 0,33 | | | 2 | | | 0,17 | | 4,70 | | | 0,0100 |
| Error | 7,57 | | | 213 | | | 0,04 | |  | | |  |
| Total | 7,91 | | | 215 | | |  | |  | | |  |
| Test: Tukey Alfa=0,05 DMS=0,07376  Error: 0,0356 gl: 213 | | | | | | | | | | | | |
| Moonlight | | | Median | | | n | | | | E.E. | | |
| Bright | | | 0,12 | | | 72 | | | | 0,02 A | | |
| Semi-bright | | | 0,15 | | | 72 | | | | 0,02 A B | | |
| Dark | | | 0,21 | | | 72 | | | | 0,02 B | | |
| ***Imantodes cenchoa.*** | | | | | | | | | | | | |
| Variance analysis | | | | | | | | | | | | |
| Variable | | N | | | R^2^ | | | R^2^ Aj | | | CV | |
| Rate | | 216 | | | 3,4 E-04 | | | 0,00 | | | 164,81 | |
| Variance analysis table (SC type III) | | | | | | | | | | | | |
| F.V. | SC | | | gl | | | CM | | F | | | p-value |
| Model | 9,7 E-04 | | | 2 | | | 4,8E-04 | | 0,04 | | | 0,9642 |
| Moonlight | 9,7 E-04 | | | 2 | | | 4,8E-04 | | 0,04 | | | 0,9642 |
| Error | 2,82 | | | 213 | | | 0,01 | |  | | |  |
| Total | 2,82 | | | 215 | | |  | |  | | |  |
| Test: Tukey Alfa=0,05 DMS=0,04503  Error: 0,0132 gl: 213 | | | | | | | | | | | | |
| Moonlight | | | Median | | | n | | | | E.E. | | |
| Bright | | | 0,07 | | | 72 | | | | 0,01 A | | |
| Dark | | | 0,07 | | | 72 | | | | 0,01 A | | |
| Semi-bright | | | 0,07 | | | 72 | | | | 0,01 A | | |
| ***Enuliophis sclateri.*** | | | | | | | | | | | | |
| Variance analysis | | | | | | | | | | | | |
| Variable | | N | | | R^2^ | | | R^2^ Aj | | | CV | |
| Rate | | 216 | | | 0,03 | | | 0,02 | | | 243,36 | |
| Variance analysis table (SC type III) | | | | | | | | | | | | |
| F.V. | SC | | | gl | | | CM | | F | | | p-value |
| Model | 0,02 | | | 2 | | | 0,01 | | 3,01 | | | 0,0512 |
| Moonlight | 0,02 | | | 2 | | | 0,01 | | 3,01 | | | 0,0512 |
| Error | 0,74 | | | 213 | | | 3,5 E-03 | |  | | |  |
| Total | 0,76 | | | 215 | | |  | |  | | |  |
| Test: Tukey Alfa=0,05 DMS=0,02302  Error: 0,0035 gl: 213 | | | | | | | | | | | | |
| Moonlight | | | Median | | | N | | | | E.E. | | |
| Semi-bright | | | 0,01 | | | 72 | | | | 0,01 A | | |
| Bright | | | 0,03 | | | 72 | | | | 0,01 A | | |
| Dark | | | 0,03 | | | 72 | | | | 0,01 A | | |
| **Other species (all data from less common species pooled)*.*** | | | | | | | | | | | | |
| Variance analysis | | | | | | | | | | | | |
| Variable | | N | | | R^2^ | | | R^2^ Aj | | | CV | |
| Rate | | 216 | | | 4,0 E-03 | | | 0,00 | | | 259,35 | |
| Variance analysis table (SC type III) | | | | | | | | | | | | |
| F.V. | SC | | | gl | | | CM | | F | | | p-value |
| Model | 0,02 | | | 2 | | | 0,01 | | 0,43 | | | 0,6499 |
| Moonlight | 0,02 | | | 2 | | | 0,01 | | 0,43 | | | 0,6499 |
| Error | 5,65 | | | 213 | | | 0,03 | |  | | |  |
| Total | 5,68 | | | 215 | | |  | |  | | |  |
| Test: Tukey Alfa=0,05 DMS=0,06373  Error: 0,0265 gl: 213 | | | | | | | | | | | | |
| Moonlight | | | Median | | | N | | | | E.E. | | |
| Semi-bright | | | 0,05 | | | 72 | | | | 0,02 A | | |
| Dark | | | 0,06 | | | 72 | | | | 0,02 A | | |
| Bright | | | 0,08 | | | 72 | | | | 0,02 A | | |

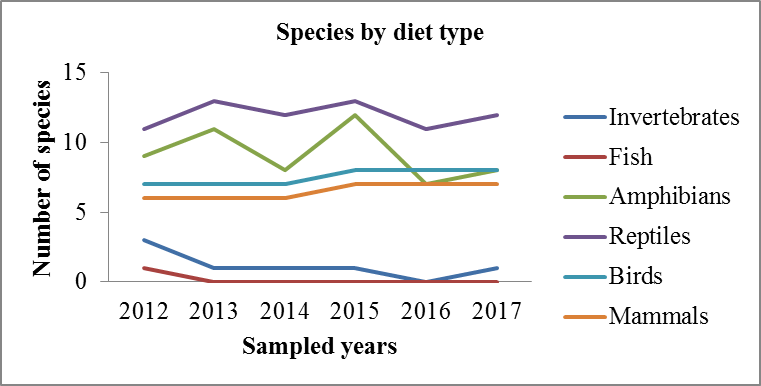

**Fig. 2**. Number of species with each diet type (diets from Savage, 2002; and Solórzano, 2004) seen per year. Species can be included on more than one category.

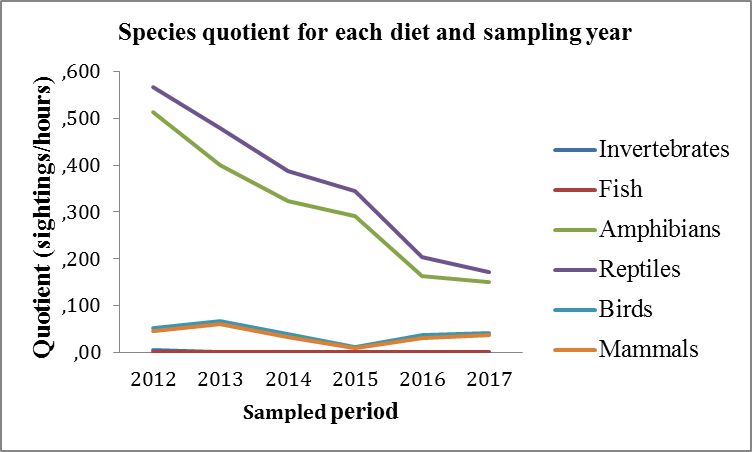

**Fig. 3**. Species counts (sightings per/hour) for each diet and sampling year

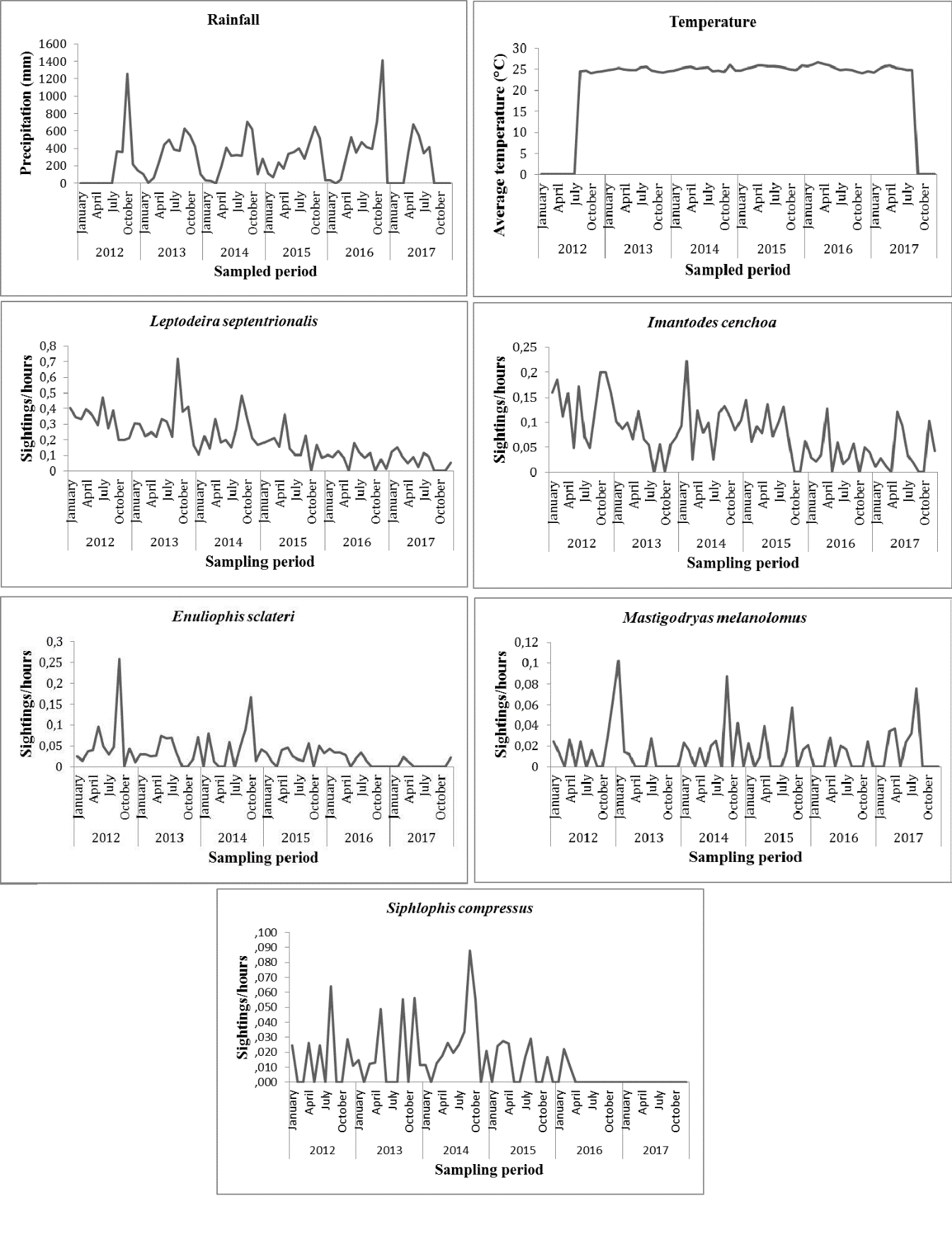

**Fig. 4.** Counts per hour of the five most frequent snakes species during the six-year study period in Drake, Costa Rica (2012-2017).

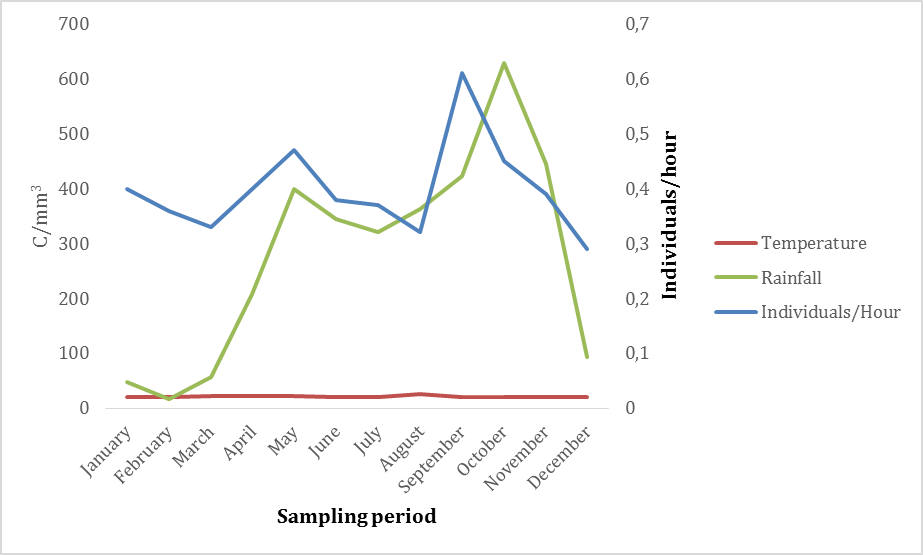

**Fig. 5.** Mean temperature, mean rainfall and individuals seen per hour for Drake snakes (Costa Rica).

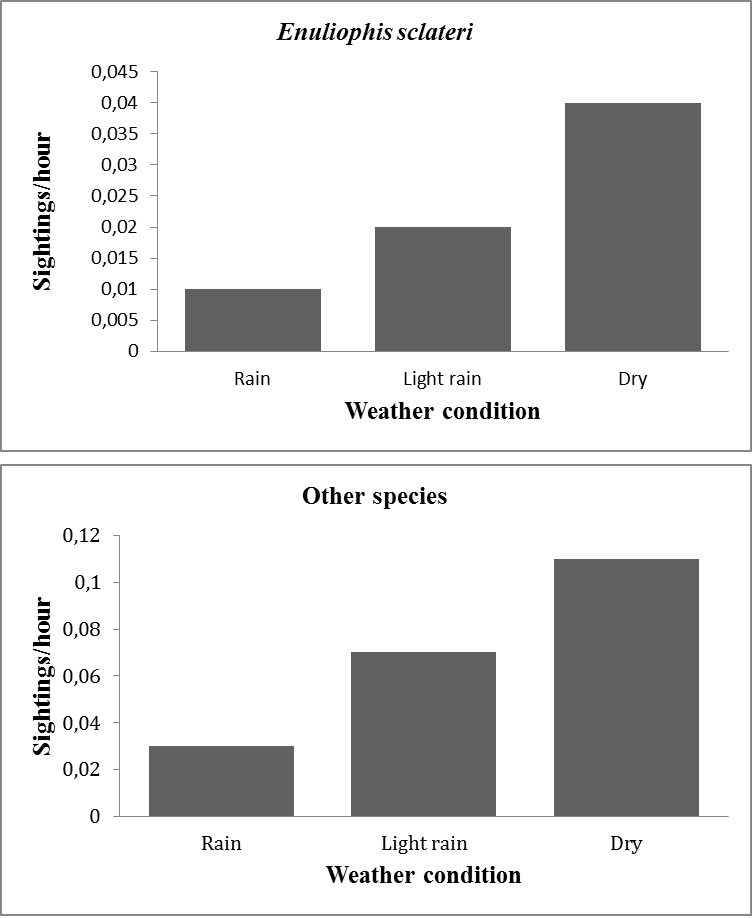

**Fig. 6.** Sightings per hour of snakes, according to rain level, in Drake, Costa Rica.

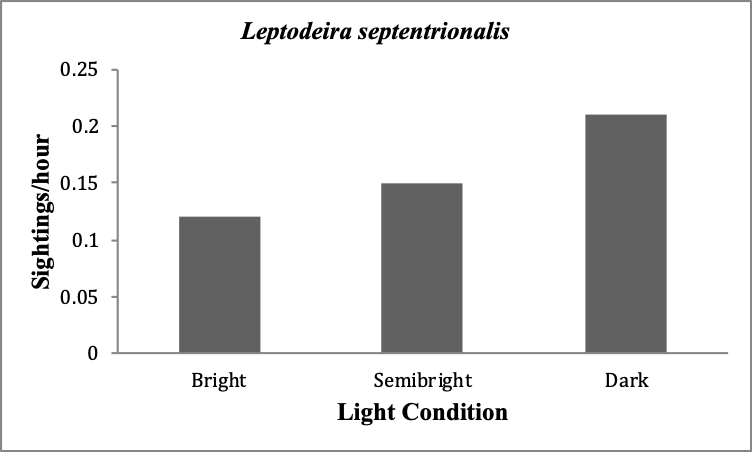

**Fig. 7.** Sightings per hour of the snake *L. septentrionalis*, according to moon light level, in Drake, Costa Rica.

*.*
