## Supplementary material for "Are tropical reptiles really declining? A six-year survey of snakes in Drake Bay, Costa Rica, and the role of temperature, rain and light": Digital appendix: DIGITAL APPENDIX 2.docx

1- Photographic catalogue of snake species found in Drake Bay, Costa Rica.

| 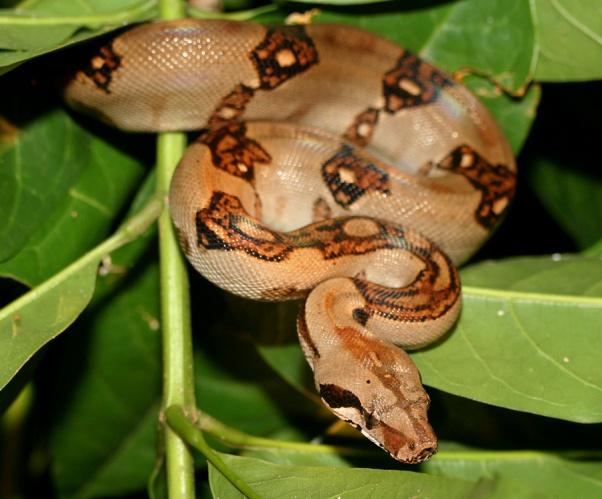  *Boa imperator* | 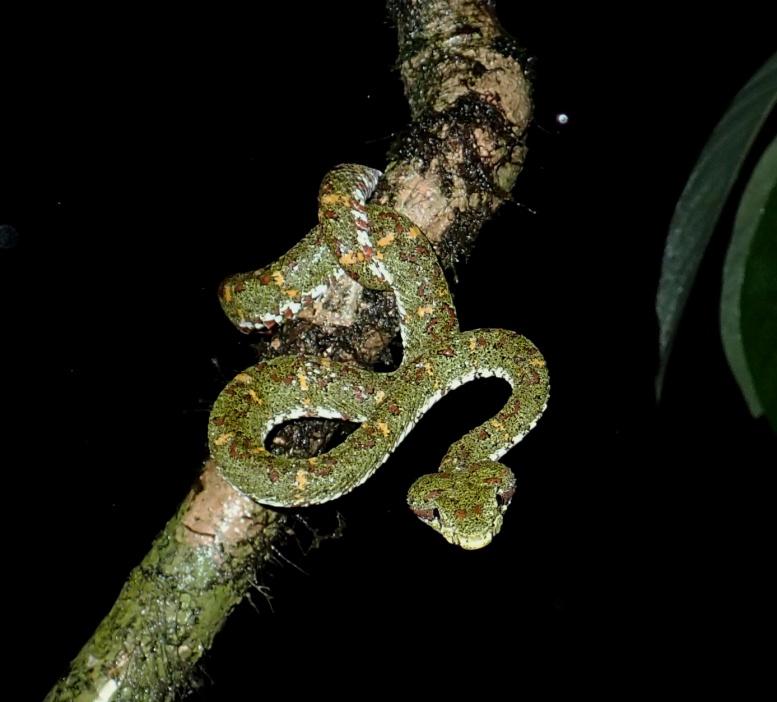  *Bothriechis schlegelii* | 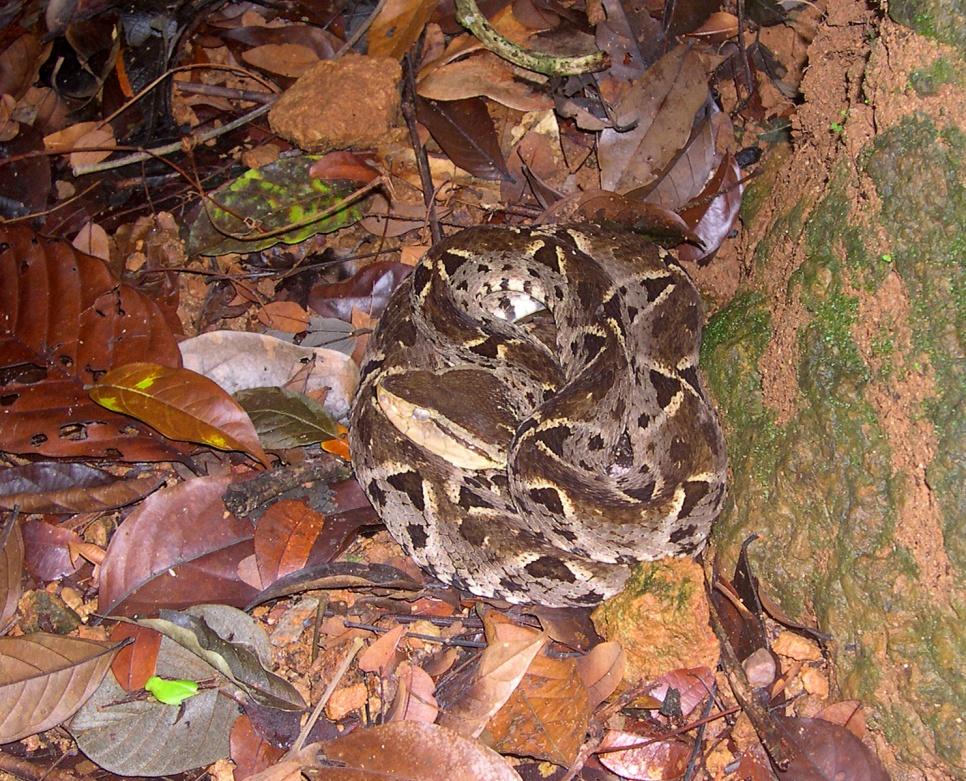  *Bothrops asper* |
| --- | --- | --- |
| 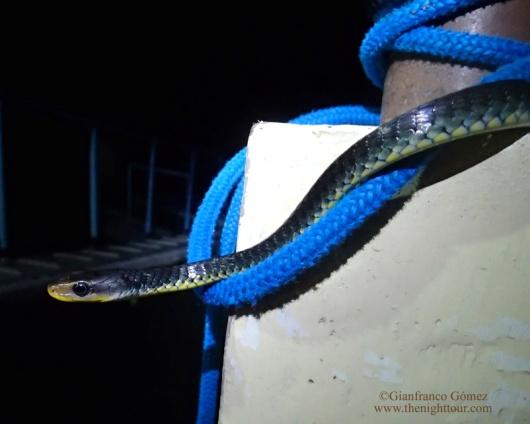  *Chironius flavopictus* | 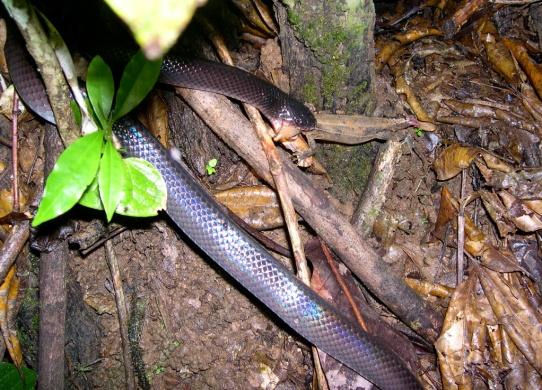  *Clelia clelia* | 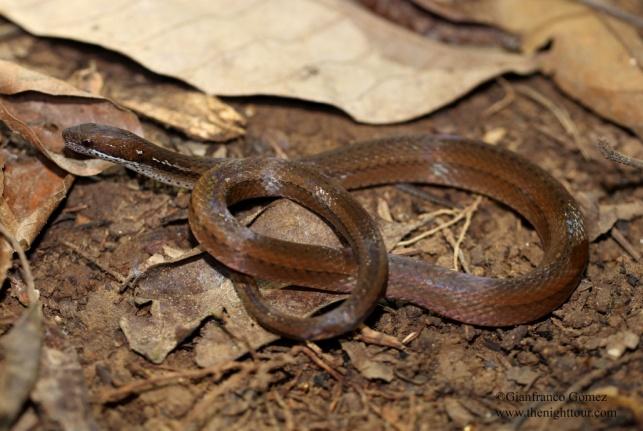  *Coniophanes fissidens* |
| 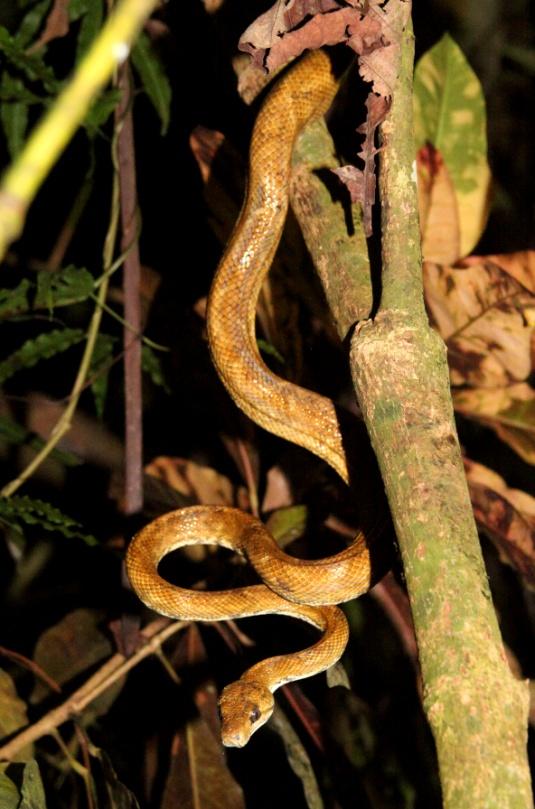  *Corallus ruschenbergerii* | 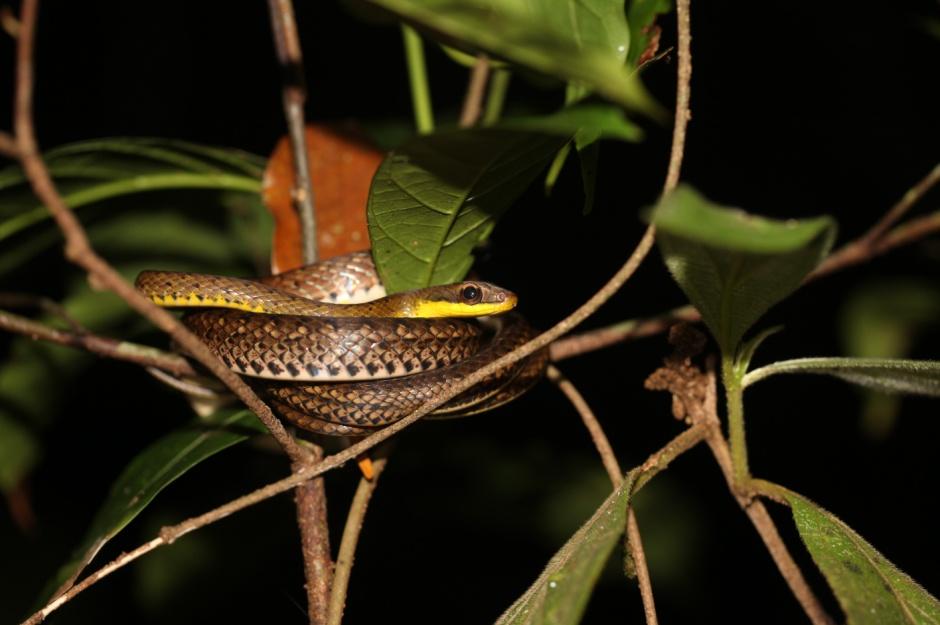  *Dendrophidion percarinatus* | 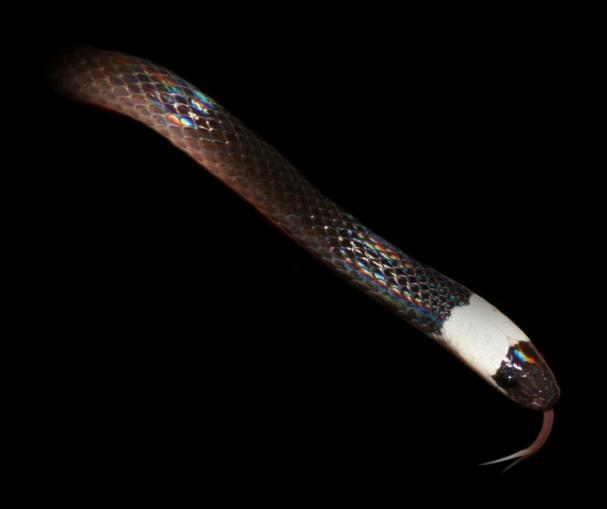  *Enuliophis sclateri* |
| 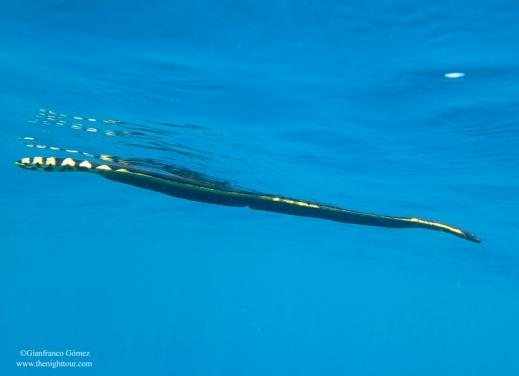  *Hydrophis platurus* | 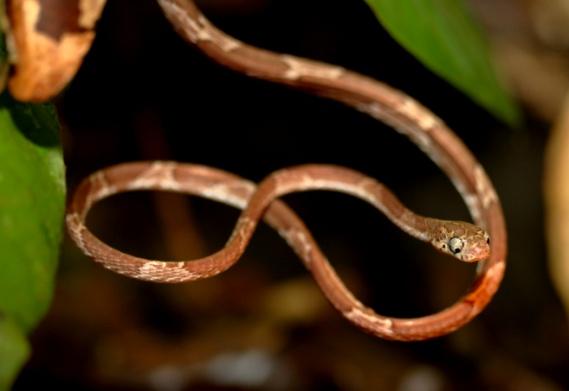  *Imantodes cenchoa* | 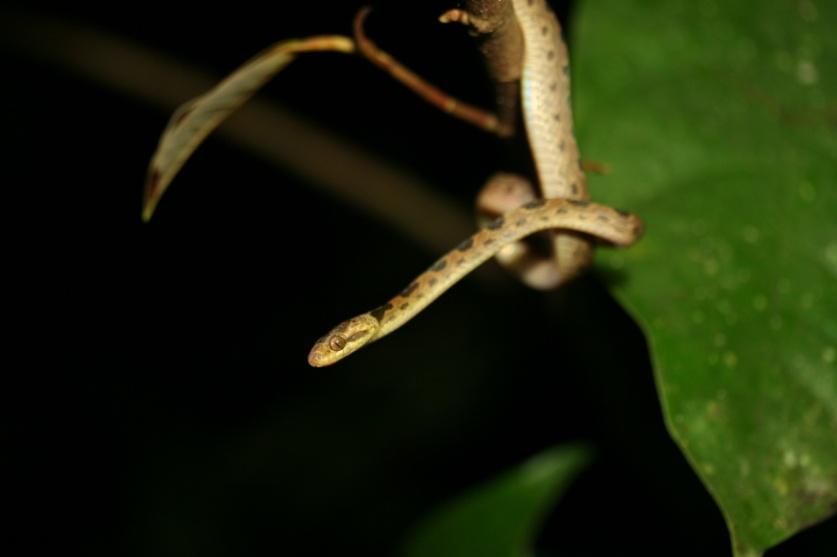  *Leptodeira septentrionalis* |
| 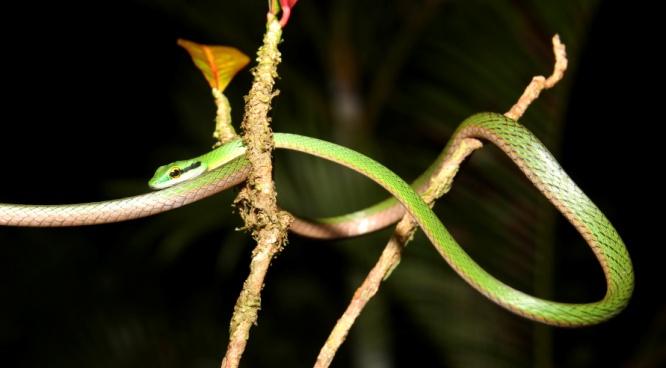  *Leptophis ahaetulla* | 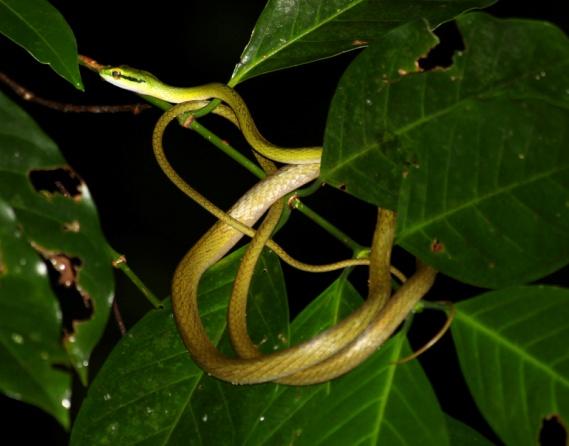  *Leptophis nebulosus* | 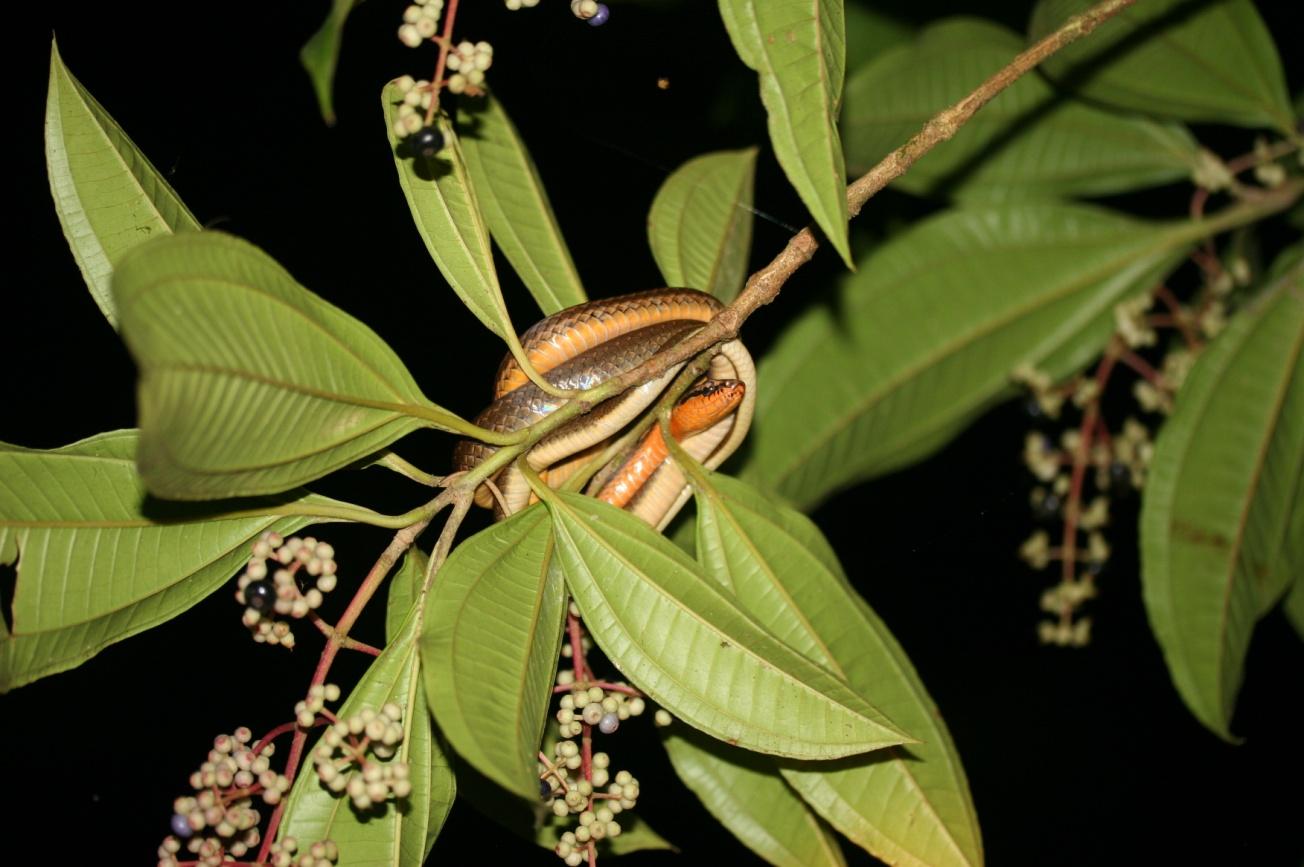  *Mastigodryas melanolomus* |
| 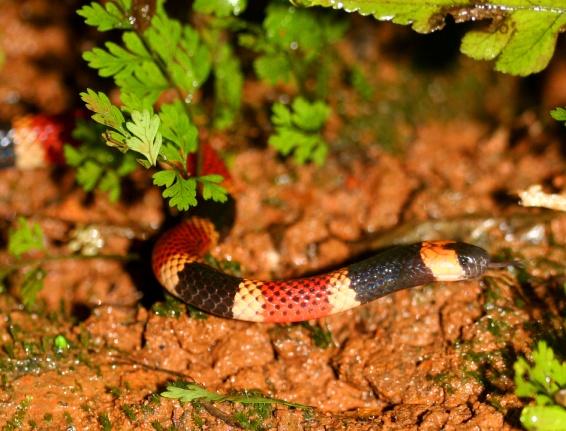  *Micrurus alleni* | 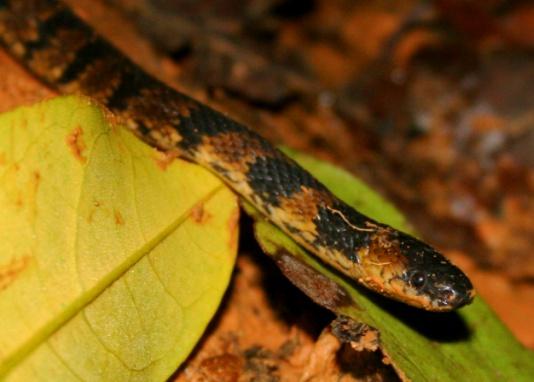  *Ninia maculata* | 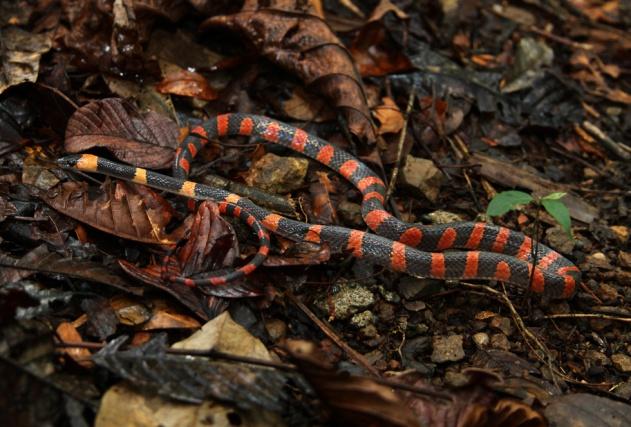  *Oxyrhopus petolarius* |
| 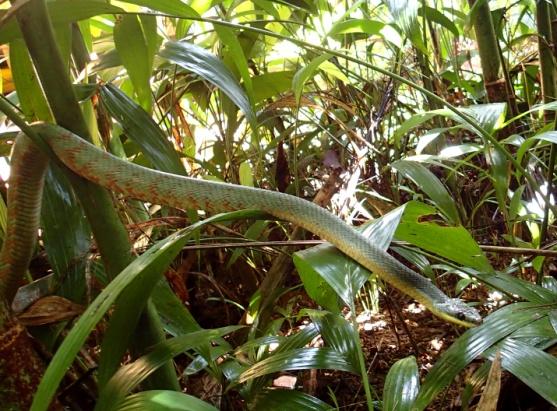  *Phrynonax poecilonotus* | 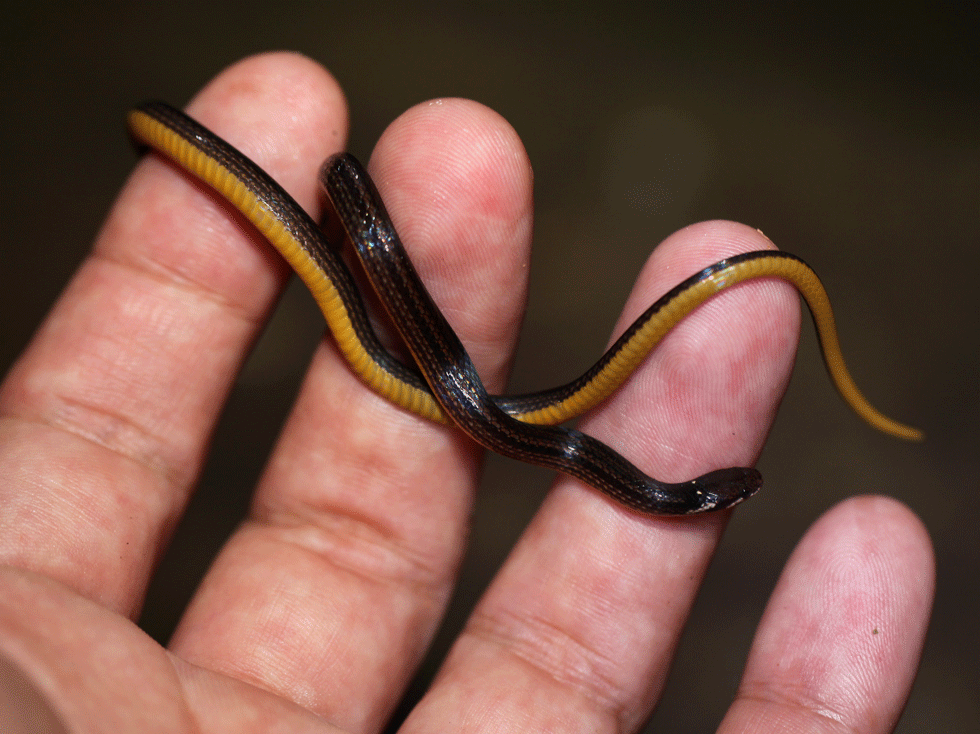  *Rhadinella godmani* | 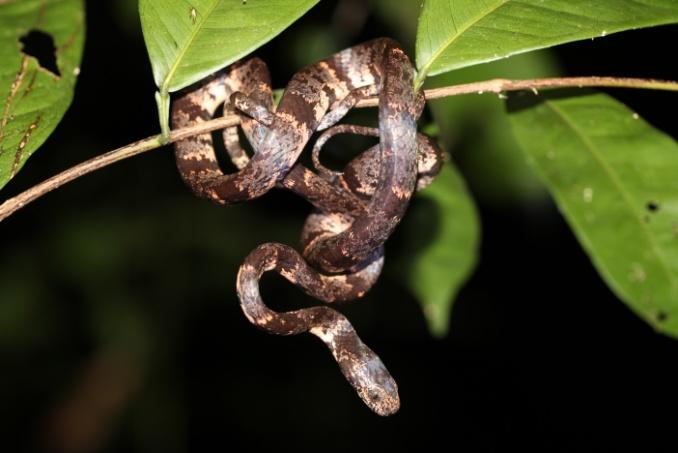  *Sibon nebulatus* |
| 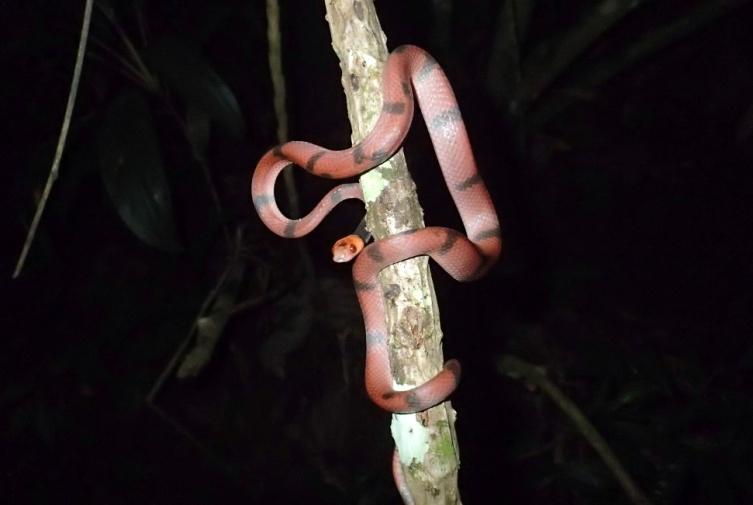  *Siphlophis compressus* | 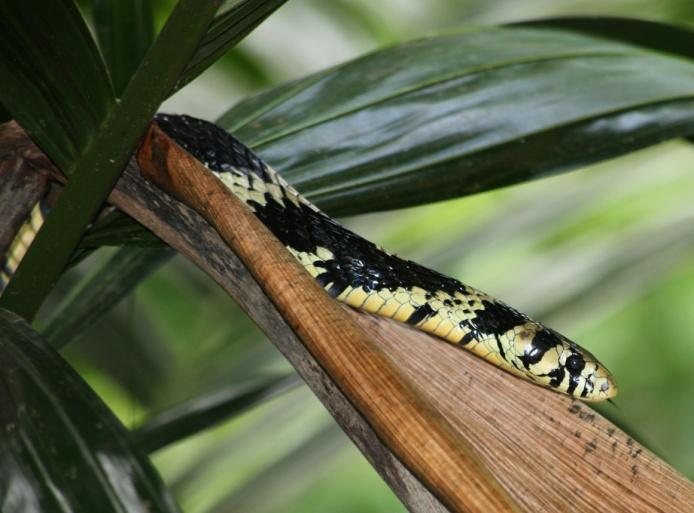  *Spilotes pullatus* | 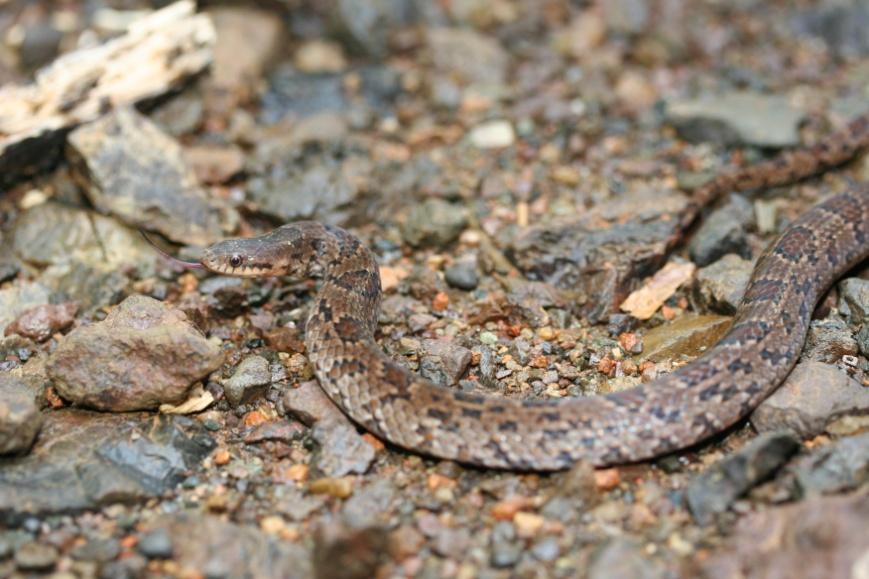  *Stenorrhina degenhardtii* |
|  | 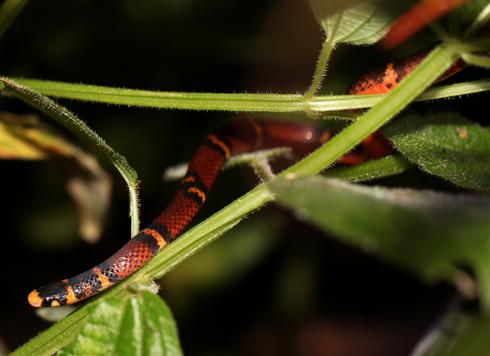  *Tantilla supracincta* |  |

2- Total sampling hours per month and year during the six year study period ion Drake, Costa Rica.

| **Month** | **Sampling hours** | | | | | |
| --- | --- | --- | --- | --- | --- | --- |
|  | **2012** | **2013** | **2014** | **2015** | **2016** | **2017** |
| **January** | 64,75 | 70,25 | 61,5 | 82,5 | 69,25 | 84,75 |
| **February** | 73,5 | 62,5 | 46 | 68,75 | 84,75 | 57,75 |
| **March** | 53,75 | 74 | 59,25 | 94,5 | 87,25 | 73,75 |
| **April** | 75,25 | 58,75 | 57,5 | 74 | 72 | 62,25 |
| **May** | 38,75 | 40,75 | 32,75 | 41,25 | 0 | 34,25 |
| **June** | 40,25 | 28 | 48,25 | 38,75 | 48,25 | 36,5 |
| **July** | 66,5 | 71 | 40 | 53,75 | 57,5 | 59,25 |
| **August** | 59,75 | 29 | 52 | 67,5 | 67,5 | 48,5 |
| **September** | 7,75 | 18,25 | 23,75 | 37,75 | 17,75 | 0 |
| **October** | 5 | 11,25 | 18 | 2,75 | 0 | 2,5 |
| **November** | 67,5 | 49,25 | 68 | 51,25 | 38,25 | 0 |

3- Sightings per hour by species, month and year in Drake Bay, Costa Rica, according to snake species.

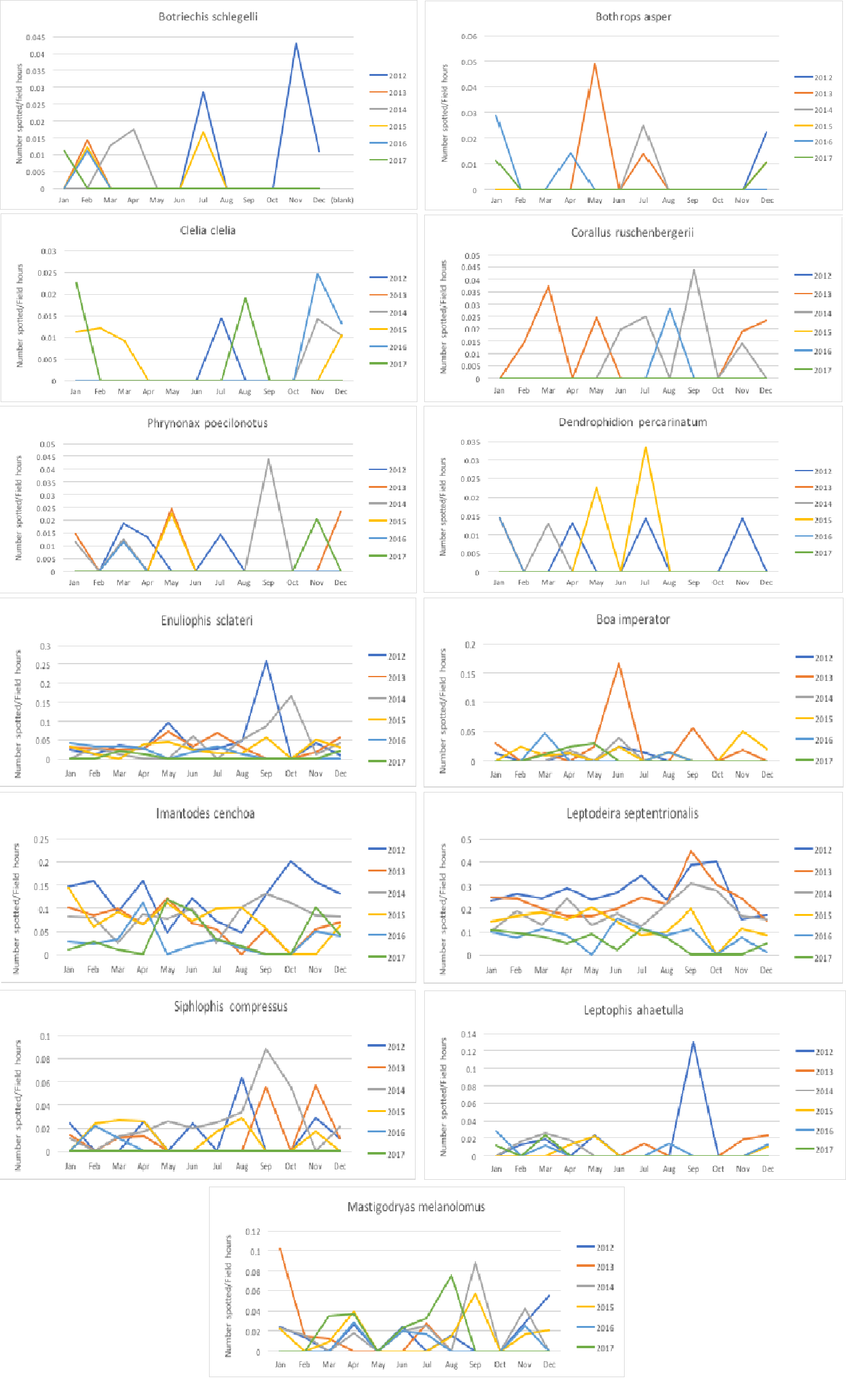

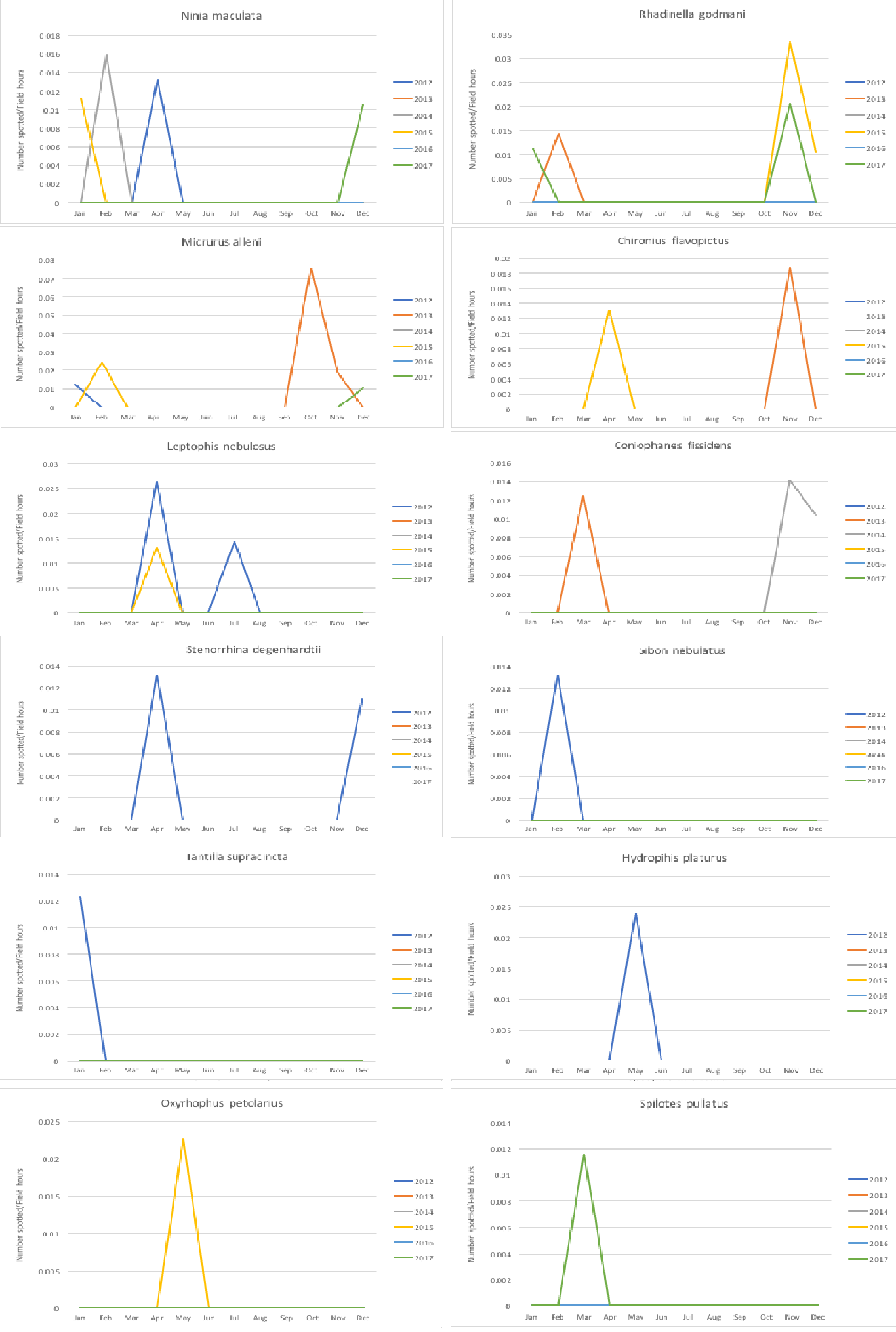

4 - Leaf litter comparison, left photograph taken on May 2013, right photograph taken on January 2018 (both by Gianfranco Gómez).

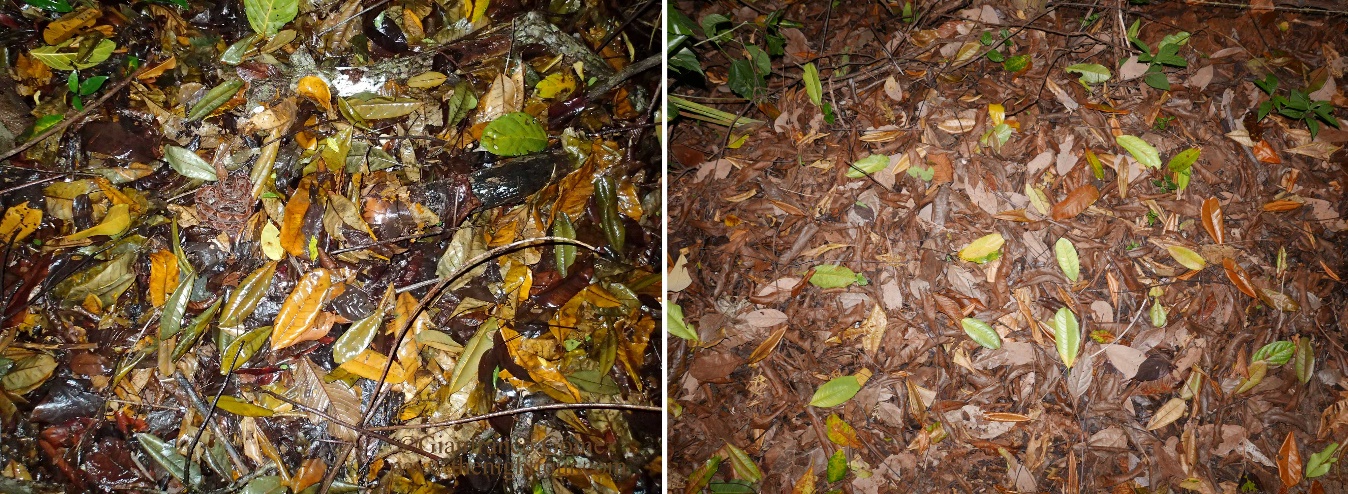
